# MCP5, a methyl-accepting chemotaxis protein regulated by both the Hk1-Rrp1 and Rrp2-RpoN-RpoS pathways, is required for the immune evasion of *Borrelia burgdorferi*

**DOI:** 10.1101/2024.06.10.598185

**Authors:** Sajith Raghunandanan, Kai Zhang, Zhang Yan, Ching Wooen Sze, Raj Priya, Yongliang Luo, Michael J Lynch, Brian R Crane, Chunhao Li, X. Frank Yang

**Author notes:** Address Correspondence to: Dr. X. Frank Yang, Dr. Chunhao Li. Equal contribution (Sajith Raghunandanan, Kai Zhang, Zhang Yan).

## Abstract

*Borrelia* (or *Borreliella*) *burgdorferi*, the causative agent of Lyme disease, is a motile and invasive zoonotic pathogen, adept at navigating between its arthropod vector and mammalian host. While motility and chemotaxis are well established as essential for its enzootic cycle, the function of methyl-accepting chemotaxis proteins (MCPs) in the infectious cycle of *B. burgdorferi* remains unclear. In this study, we demonstrate that MCP5, one of the most abundant MCPs in *B. burgdorferi*, is differentially expressed in response to environmental signals as well as at different stages of the pathogen’s enzootic cycle. Specifically, the expression of *mcp5* is regulated by the Hk1-Rrp1 and Rrp2-RpoN-RpoS pathways, which are critical for the spirochete’s colonization of the tick vector and mammalian host, respectively. Infection experiments with an *mcp5* mutant revealed that spirochetes lacking MCP5 could not establish infections in either C3H/HeN mice or Severe Combined Immunodeficiency (SCID) mice, which are defective in adaptive immunity, indicating the essential role of MCP5 in mammalian infection. However, the *mcp5* mutant could establish infection and disseminate in NOD SCID Gamma (NSG) mice, which are deficient in both adaptive and most innate immune responses, suggesting a crucial role of MCP5 in evading host innate immunity. In the tick vector, the *mcp5* mutants survived feeding but failed to transmit to mice, highlighting the importance of MCP5 in transmission. Our findings reveal that MCP5, regulated by the Rrp1 and Rrp2 pathways, is critical for the establishment of infection in mammalian hosts by evading host innate immunity and is important for the transmission of spirochetes from ticks to mammalian hosts, underscoring its potential as a target for intervention strategies.

**SUMMARY:** Lyme disease is the most commonly reported arthropod-borne illness in the US, Europe, and Asia. The causative agent of Lyme disease, *Borrelia burgdorferi*, is maintained in an enzootic cycle involving arthropod vectors (*Ixodes* ticks) and rodent mammalian hosts. Understanding how *B. burgdorferi* moves within this natural cycle is crucial for developing new strategies to combat Lyme disease. The complex nature of the enzootic cycle necessitates sensory-guided movement in response to environmental stimuli. *B. burgdorferi* possesses a unique and intricate chemotaxis signaling system, with methyl-accepting chemotaxis proteins (MCPs) at its core. These proteins are responsible for sensing environmental signals and guiding bacterial movement toward or away from stimuli. This study found that one of the MCPs, MCP5, is highly expressed and differentially regulated during the enzootic cycle by the Hk1-Rrp1 and Rrp2-RpoN-RpoS pathways. MCP5 is crucial for mammalian infection, aiding in immune evasion and transmission from ticks to mammals, providing a foundation for further research into *B. burgdorferi*’s navigation within its hosts.

## INTRODUCTION

Chemotaxis allows motile bacteria to swim towards a favorable environment or away from one that is toxic, which has been well characterized in the two paradigm model organisms *Escherichia coli* and *Salmonella enterica* Typhimurium [1–3]. Bacterial chemotaxis is modulated through a signaling cascade that are composed of chemoreceptors, a coupling protein CheW, a histidine kinase CheA, and a response regulator CheY [3–5]. Bacterial chemoreceptors, also known as <u>m</u>ethyl-accepting <u>c</u>hemotaxis <u>p</u>roteins (MCPs), typically contain four functional units, including a periplasmic ligand-binding domain, a transmembrane region, a cytoplasmic HAMP (<u>h</u>istidine kinase, <u>a</u>denylyl cyclase, <u>m</u>ethyl-accepting chemotaxis protein, and <u>p</u>hosphatase) domain and a kinase-control module [6]. MCPs form trimers of dimers in an array-like structure that typically resides at bacterial cell poles and sense a variety of ligands (e.g., attractants or repellents) [7, 8]. Ligand binding to the MCPs, either alone or together with one of the periplasmic binding proteins, promotes a conformational change in the receptor that modulates the activity of CheA. Activated CheA transfers a phosphoryl group to CheY, generating phosphorylated CheY (CheY-P) which in turn interacts with the motor switch complex (also known as C-ring) to control flagellar rotation and locomotion [6]. When the attractant concentration remains stable, bacteria adapt through a process that involves methylation of glutamate residues in the cytoplasmic domains of MCPs [8]. In addition to chemotaxis, MCPs are also implicated in the regulation of biofilm formation [9], flagellum biosynthesis [10], degradation of xenobiotic compounds [11], and production of toxins [12].

*Borrelia* (or *Borreliella*) *burgdorferi*, the causative agent of Lyme disease, is a motile and invasive spirochetal pathogen [13, 14]. Motility and chemotaxis are critical for spirochetes to be maintained in the enzootic cycle between tick vectors and vertebrate hosts. When ticks acquire spirochetes from infected vertebrate hosts upon blood feeding, spirochetes are attracted to the tick feeding site by chemotactic signals [15, 16]. When infected ticks transmit *B. burgdorferi* to naïve vertebrate hosts via feeding, spirochetes replicate, exit the tick gut, move to tick hemocoel, and then migrate to the salivary gland, and are subsequently transmitted to vertebrate hosts [17]. Within vertebrate hosts, *B. burgdorferi* cells disseminate from the infection site, and are capable of penetrating host connective tissue and invading various organs such as joints, heart, and nervous system, and causing multi-stage diseases [14, 18, 19]. In line with its important role in the enzootic cycle, *B. burgdorferi* has a unique and complex chemotaxis signaling system [20]. Its genome encodes multiple copies of chemotaxis genes, including two histidine kinases (CheA1 and CheA2), three response regulators (CheY1, CheY2, and CheY3), three coupling proteins (CheW1, CheW2, and CheW3), two sets of chemotaxis adaptation proteins, CheB (CheB1 and CheB2) and CheR (CheR1 andCheR2), and five MCPs (MCP1, MCP2, MCP3, MCP4, and MCP5) and one cytoplasmic chemoreceptor [20–26]. Several motility mutants (e.g., *fliG1, flaB, fliH, fliL, flhF,* and *motB*) and chemotaxis mutants (e.g., *cheA1*, *cheA2, cheY1, cheY2, cheY3,cheX*, and *cheD*) were reported, and the results demonstrated that motility and chemotaxis are essential for spirochetes to survive and colonize in both ticks and vertebrate hosts [for recent review, see [27]]

How *B. burgdorferi* regulates motility and chemotaxis during its enzootic cycle has not been elucidated. In this regard, several regulators and signaling pathways have been identified to coordinately regulate differential gene expression of *B. burgdorferi* during the infection [for review, see recent reviews [28, 29]]. Among these, Hk1-Rrp1 and Rrp2-RpoN-RpoS pathways play central roles in controlling differential expression of genes critical for tick colonization and mammalian infection, respectively [14, 30–32]. The Hk1-Rrp1 two-component signaling pathway senses unknown signals and becomes activated, resulting in the production of a second messenger c-di-GMP [33–35]. This pathway is required for *B. burgdorferi* to survive in feeding ticks and complete the enzootic life cycle [33–35]. Hk1-Rrp1 controls the expression of genes important for spirochetal utilization of glycerol, chitobiose, and N-acetylglucosamine, as well as for the process of chemotaxis, motility, and osmolality sensing [33, 34, 36–41]. On the other hand, the Rrp2-RpoN-RpoS pathway, also called σ^N^-σ^S^ alternative σ factor cascade, is activated by Rrp2 and RpoN (σ^N^) when spirochetes transmit to the mammalian host and during the phase of mammalian infection, resulting in the production of alternative sigma factor RpoS (σ^S^) [42–46]. RpoS, as a global regulator, further activates the transcription of many virulence genes essential for transmission and infectivity in vertebrate hosts, while repressing the expression of genes required for spirochete survival in the tick vector [14, 28, 42, 43].

Compared to other chemotaxis proteins, little is known about the function of MCP chemoreceptors in *B. burgdorferi*. Although several chemoattractants of *B. burgdorferi* have been identified [15, 47–49], the MCP proteins responsible for sensing these attractants remain unknown. The lack of knowledge regarding MCP function has hindered our understanding of how *B. burgdorferi* navigates between and within the tick vector and the vertebrate host. In this study, we concentrated on one of the highly expressed MCPs, MCP5, and discovered that its expression is differentially regulated during the enzootic cycle of *B. burgdorferi*, controlled by both the Hk1-Rrp1 pathway and the Rrp2-RpoN-RpoS pathway. We further demonstrate that MCP5 plays a pivotal role in mammalian infection by aiding spirochetal evasion of the host’s innate immune response, as well as contributing to spirochetal transmission from ticks to mammals.

## RESULTS

### Structural analyses of *B. burgdorferi* MCPs

The genome of *B. burgdorferi* encodes five putative chemoreceptors, including MCP1 (BB0578), MCP2 (BB0596), MCP3 (BB0597), MCP4 (BB0680), and MCP5 (BB0681). Our previous study reveals that these MCP proteins form a long, thin array-like structure that resides at the cell poles of *B. burgdorferi* [50]; however, their roles in chemotaxis remain largely unknown. To address this question, we first constructed their homology structures using AlphaFold and then compared these structures to their counterparts from other bacteria. Overall, MCPs 2-5 share similar domain composition and structural topology, with MCP1 being the most distant (**Fig. 1**). Unlike other MCPs, MCP1 is short (∼21.6 nm) and has no N-terminal periplasmic ligand-binding domain. Instead, it has a C-terminal ligand binding domain (**Fig. 1A**), suggesting it may function as a cytoplasmic MCP that senses internal signals. MCPs 2-4 form long helical structures with different lengths, ranging from 272 A to 452 A (**Fig. 1B-E**). Among these five MCPs, MCP3 is the longest (452 A, **Fig. 1C**) because it contains several specific inserts. Multiple sequence alignments further revealed that MCPs 1-5 possess a conserved protein-interaction region (PIR) interacting with CheA/CheW (**Fig. S1**). Collectively, these results indicate that MCPs 1-5 are canonical chemoreceptors albeit with some sequence and structural variations.

**Fig. 1.**
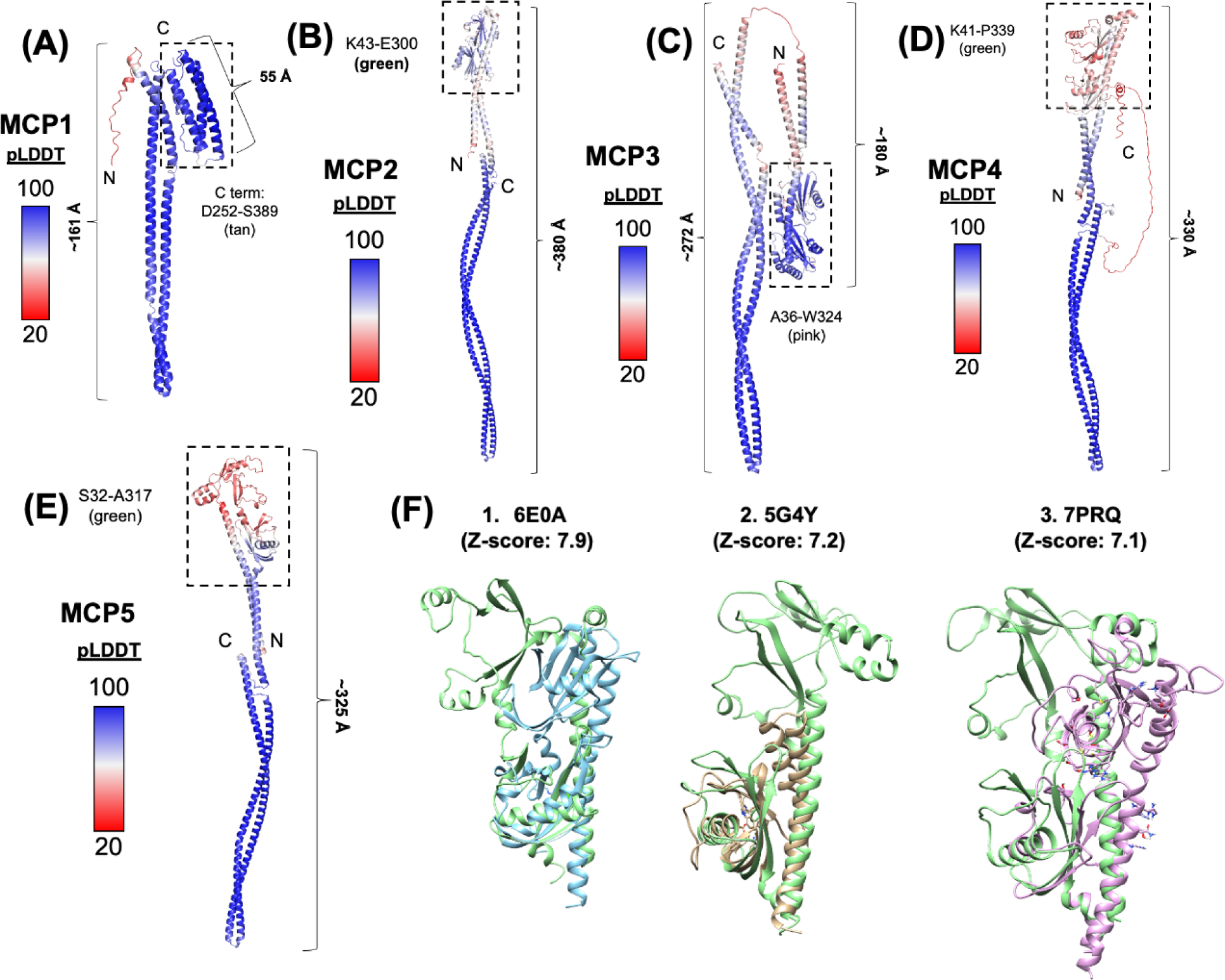
Structural Analysis of MCP1-5 of *B. burgdorferi*. AlphaFold models of **(A)** MCP1, **(B)** MCP2, **(C)** MCP3, **(D)** MCP4 and **(E)** MCP5. Protein structures are colored according to their pLDDT scores [51]. For each MCP, the sensor domain is marked with dotted black box. **(F)** Structural comparison of the Bb MCP5 sensor domain against similar proteins structures in the PDB. The top three hits were the ligand-binding domain of *Helicobacter pylori* chemoreceptor TlpA (PDB: 6E0A)[52], the cache-like sensor domain PscD-SD from the plant pathogen *Pseudomonas syringae* (PDB: 5G4Y)[53] and the ligand binding domain of the chemoreceptor PctD from *Pseudomonas aeruginosa* (PDB: 7PRQ)[54]. Structures were identified using the DAHLI server [55].

### *mcp5* is highly expressed *in vitro*

Using qRT-PCR, we examined the expression levels of five *mcp* genes in *B. burgdorferi*. The result showed that *mcp4* and *mcp5* are the two most highly expressed genes when spirochetes were cultured under *in vitro* growth conditions (**Fig. 2A**). *mcp4* and *mcp5* are adjacent to each other in the *B. burgdorferi* genome (**Fig. 2B**) [26]. 5′RACE analysis revealed that the transcriptional start sites (TSS) of *mcp4* and *mcp5* are at the same position (G), 62 bp upstream from the ATG start codon of *mcp4* (**Fig. 2B, C**), indicating these two genes are co-transcribed by the same promoter. A putative -10/-35 σ^70^-like promoter sequence was also identified 6 bp upstream of TSS (**Fig. 2B**). Given that *mcp5* is highly expressed and it is located at the end of the *mcp4*-*mcp5* operon, we focused on *mcp5* in this study.

**Fig. 2.**
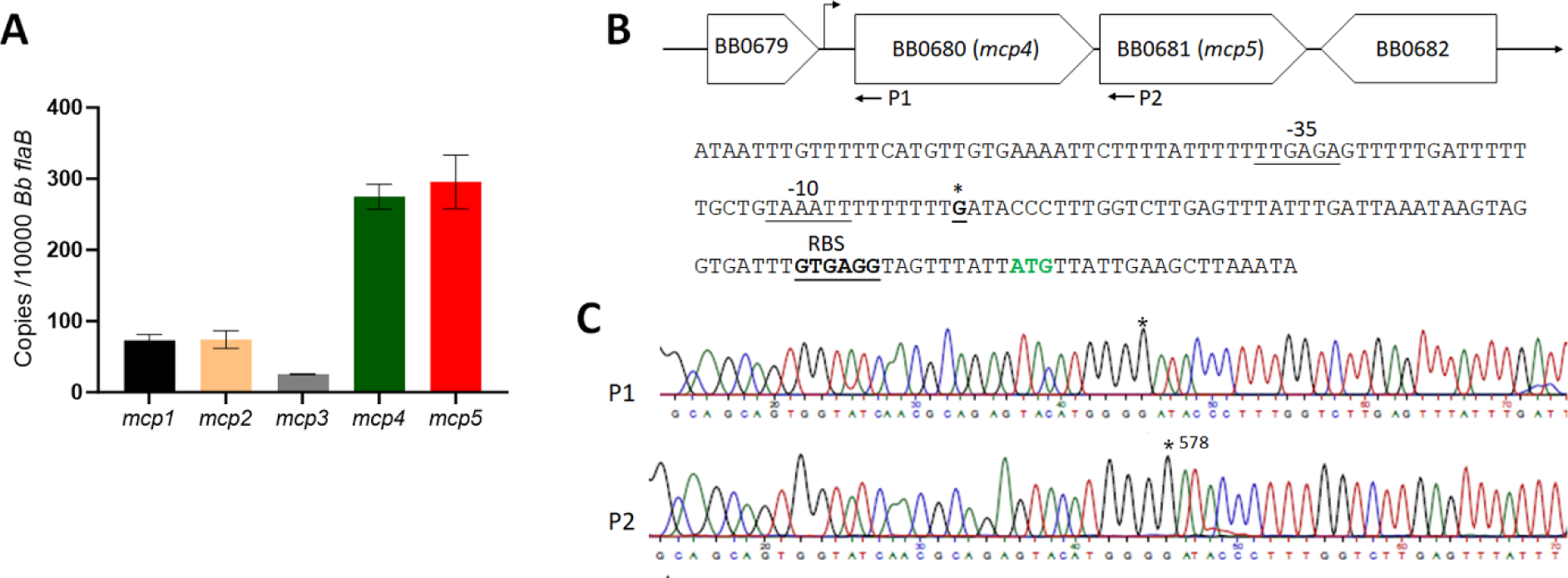
Expression of five *mcp* genes and identification of *mcp5* promoter. (**A**) **qRT-PCR analyses.** Wild-type *B. burgdorferi* strain B31M was cultured in the BSK-II medium at 37°C and harvested at the mid-log phase. RNAs were extracted and subjected to qRT-PCR analyses to examine the expressions of all five *mcp* genes. Levels of *mcp* expression relative to the level of *flaB* expression were reported. The bars represent the mean values of three independent experiments. (**B**) Schematic presentation of *mcp4* and *mcp5* in the *B. burgdorferi* genome (top) and the promoter sequence at the flanking region of *mcp4* (bottom). The asterisk (*) indicates the transcription start site. The predicted -35 and -10 promoter and RBS sequences are underlined and labeled. ATG in green denotes the start codon of *mcp4*. (**C**) 5′RACE analysis illustrated with a DNA sequencing chromatogram, where the asterisk (*) indicates the +1 position.

### *mcp5* expression is regulated by environmental cues

Many virulence genes of *B. burgdorferi* are differentially expressed in the enzootic cycle and regulated by environmental cues, such as temperature, pH, and cell density[56–58]. To investigate whether *mcp5* expression is influenced by culture temperature, pH or cell density, spirochetes were under different culture conditions or harvested at different growth phases, such as mid-log (M) vs stationary (S) growth phases. The resulted samples were subjected to qRT-PCR and immunoblotting analyses. The result showed that the expression of *mcp5* was induced by higher cell density, temperature, and lower pH (37°C, pH 7.0) (**Fig. 3**), a condition mimicking tick feeding conditions [56–58]. The observed expression pattern of *mcp5* is similar to that of *ospC*, suggesting that they are probably controlled by same regulators.

**Fig. 3.**
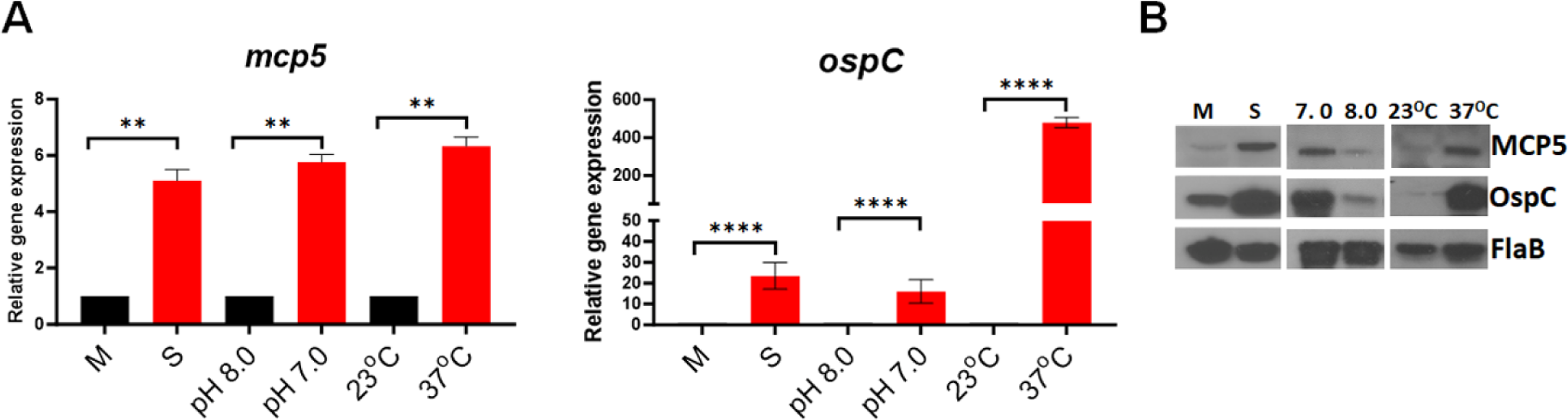
Influence of *mcp5* expression by various environmental cues. (**A**) qRT-PCR analyses. Wild-type *B. burgdorferi* strain B31M were cultures in standard BSK-II medium (pH 7.5) at 37°C and harvested at different growth phases [mid-log (M) or stationary (S)], cultured in BSK-II medium under different pH (8.0 vs 7.0), or different temperatures (23°C vs 37°C). RNAs were extracted and subjected to qRT-PCR analyses for the expressions of *mcp5* (left) and *ospC* (right). The relative gene expressions are recorded, with the levels of gene expression in cultures M, pH 8.0, and 23°C normalized to 1.0. The bars represent the mean values of three independent experiments, and the error bars represent the standard deviation. ****, *p* < 0.00001, ** *p* < 0.001 respectively using one-way ANOVA. (**B**) Western blot analyses of the whole cell lysates of spirochetes from (**A**) and (**B**). FlaB was used as a loading control.

### *mcp5* expression is induced during tick feeding and mammalian infection

To investigate the expression of *mcp5* in the enzootic cycle of *B. burgdorferi*, pathogen-free, unfed *I. scapularis* larvae were fed on mice (C3H/HeN) infected with wild-type *B. burgdorferi* strain B31M. Fed larvae were allowed to molt to the nymphal stage. Infected flat nymphs were then allowed to feed on naive mice for transmission. Feeding nymphs were collected at 48 or 96 hrs post-feeding. As shown in **Fig. 4A**, the *mcp5* transcripts were undetectable in flat nymphs, and blood feeding induced *mcp5* expression at 48 hrs and further induced at 96 hours of feeding. This result indicates that *mcp5* expression is induced upon blood feeding in the transmission phase of the cycle.

**Fig. 4.**
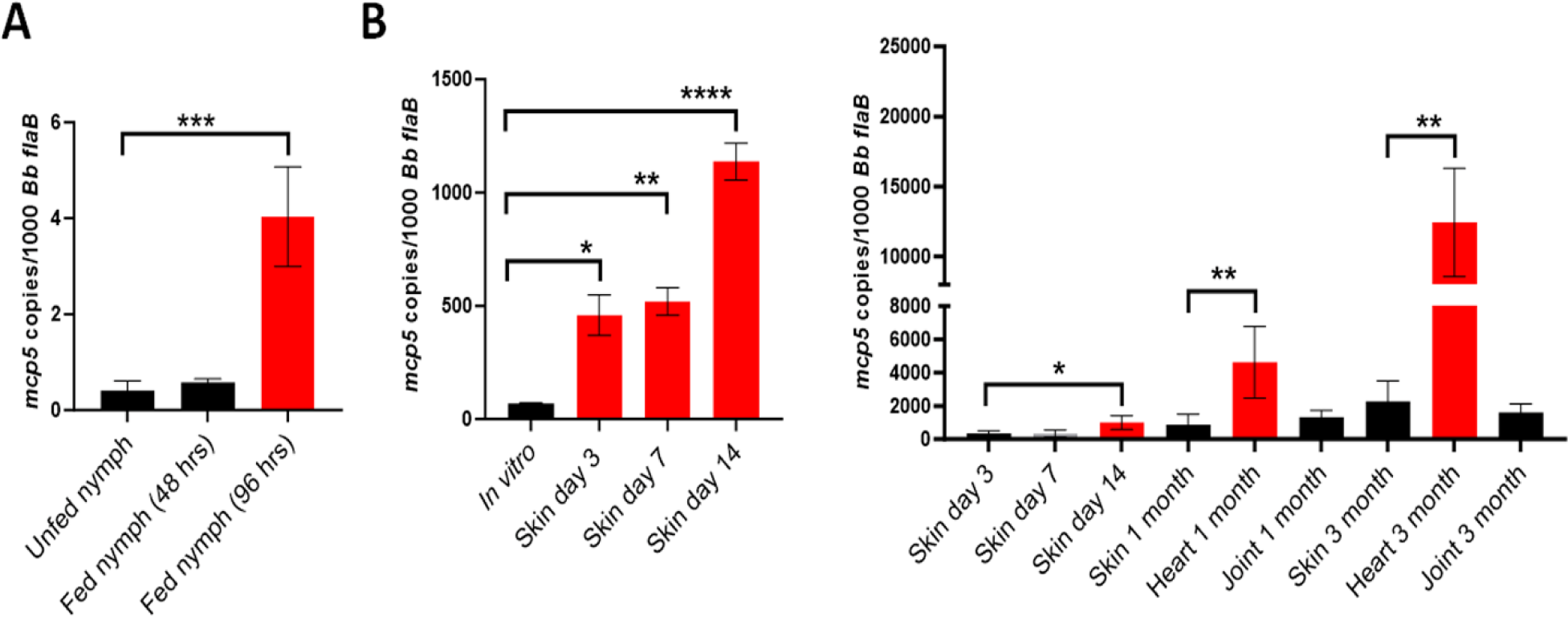
Expression of *mcp5* in the enzootic cycle of *B. burgdorferi*. Relative *mcp5* expression in various stages in ticks (**A**) and mice (**B**) was determined by qRT-PCR analyses and reported as copies of *mcp5* per 1,000 copies of the *flaB* transcripts. For gene expression in ticks, infected unfed nymphs or fed nymphs (48- and 96-hours post-feeding) containing *B. burgdorferi* were determined by qRT-PCR analyses. The bar represents values from 10 data points, and each data point was generated from 10 unfed nymphs or 1 fed nymph. For mouse experiments, mice were euthanized on days 3, 7, 14, 30 (1 month) and 90 (3 month) post-infection, and skin (site of infection), heart and joint tissues were collected and subjected to RNA extraction followed by qRT-PCR analyses. The bar represents the average values of *mcp5* transcripts calculated from 5 independently infected mice performed in triplicates. The error bars represent the standard deviation. *, *p* < 0.01; **, *p* < 0.001; ***, *p* < 0.0001 using one-way ANOVA.

To determine the *mcp5* expression the mammalian host, mice were sacrificed at various time points after the infection and skin, joint and heart tissues were collected and subjected to qRT-PCR analyses. The result showed that relative to its expressions in ticks and *in vitro*, the *mcp5* expression had a much higher level of expression in mice (**Fig. 4B**). This suggests that *mcp5* expression is further induced when spirochetes replicate in the mammalian host. Interestingly, the *mcp5* expression showed significantly higher levels in heart tissues (1 and 3 months of post infection) than that in other tissues (**Fig. 4B**, right panel).

### *mcp5* is regulated by both the Hk1-Rrp1 and Rrp2-RpoN-RpoS pathways

Given that the Hk1-Rrp1 pathway and the Rrp2-RpoN-RpoS pathway are the two important pathways that control differential expression of many genes essential for colonization in ticks or infection in mammals [14, 30–32], we sought to investigate whether *mcp5* is regulated by these two pathways. To this end, we measured the expression of *mcp5* in various mutants that are defective in these two pathways using qRT-PCR. Our results showed that level of *mcp5* expression was significantly downregulated in all the mutants but restored to the wild-type level in their isogenic complemented strains (**Fig. 5A**). In consistent with this finding, immunoblot results showed that all the mutants had diminished level of MCP5 (**Fig. 5B**). These results suggest that the *mcp5* expression is controlled by both the Hk1-Rrp1 and Rrp2-RpoN-RpoS pathways.

**Fig. 5.**
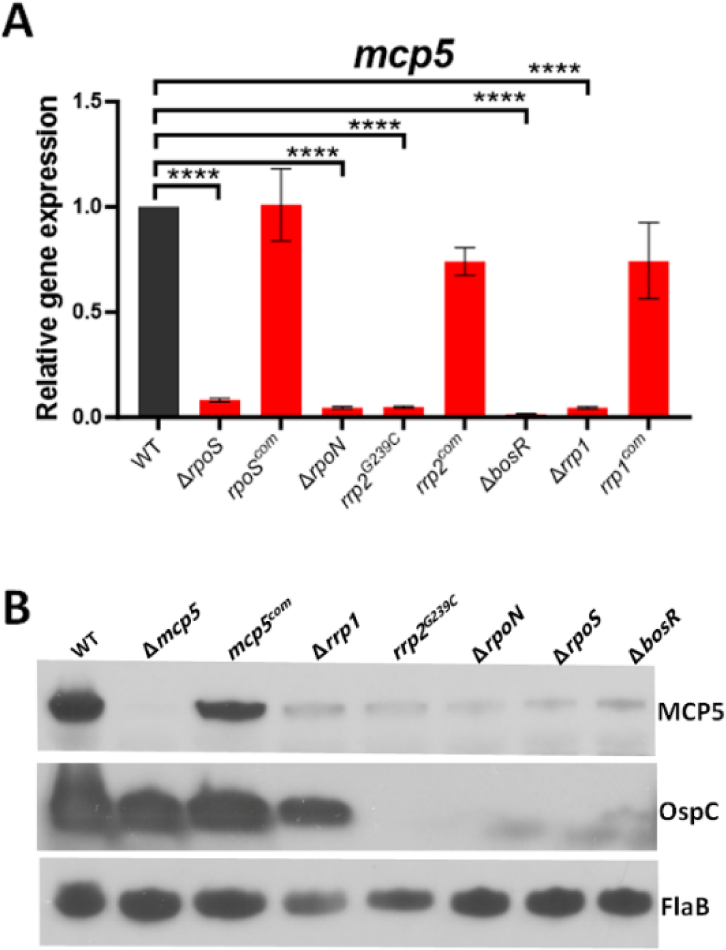
Influences of *mcp5* expression by Rrp1-Hk1 and Rrp2-RpoN-RpoS pathways. **(A)** qRT-PCR analyses. Wild-type *B. burgdorferi* strain B31 (WT), *rpoS* mutant ( Δ *rpoS*) and its isogenic complemented strain (*rpoS^com^*), *rrp2* point mutant (*rrp2^G239C^*) and complemented strain (*rrp2^com^*), *bosR* mutant (Δ*bosR*), *rrp1* mutant (Δ*rrp1*) and complemented strain (*rrp1^com^*) were cultured in BSK-II medium at 37°C and harvested at late-log phase. RNAs were extracted and subjected to qRT-PCR analyses for *mcp5* expressions. The levels of *mcp5* expression in all strains were first normalized with the level of *flaB*. Then, relative levels of *mcp5* expression to the levels in wild-type *B. burgdorferi* (which were normalized to 1.0) are reported. The bars represent the mean values of three independent experiments, and the error bars represent the standard deviation. ****, *p* < 0.00001, * *p* < 0.01 respectively using one-way ANOVA. (**B**) Immunoblot analyses. Whole cell lysates of various *B. burgdorferi* strains were harvested and subjected to immunoblotting using antibodies against MCP5, OspC, or FlaB (loading control).

### Construction of a *mcp5* mutant and its isogenic complemented strain

To investigate the role of MCP5 in the enzootic cycle of *B. burgdorferi*, a *mcp5* mutant and its complemented strain were constructed as illustrated in **Fig. 6A**. The loss and restoration of MCP5 production in the *mcp5* mutant (Δ*mcp5*) and complemented strain (*mcp5^com^*) were confirmed by PCR (**Fig. 6B**) and immunoblotting (**Fig. 6C**). Deletion of *mcp5* did not affect the production of MCP4 whose gene was located upstream of *mcp5* (**Fig. 6C**), indicating that deletion of *mcp5* has no polar effect on *mcp4* expression. Subsequent endogenous plasmid profile analyses showed that the *mcp5* mutant and its complemented strain lost cp32-6 and lp28-4 (**Fig. 6D**). Since these two plasmids are not required for infectivity [59], we proceeded phenotypical characterizations with these two strains. Deletion of *mcp5* had no impact on *B. burgdorferi* growth, swimming behaviors, and its response to N-acetylglucosamine (NAG) and rabbit serum, two chemoattractants of *B. burgdorferi* (Figures S2 and 3) [47, 48].

**FIG. 6.**
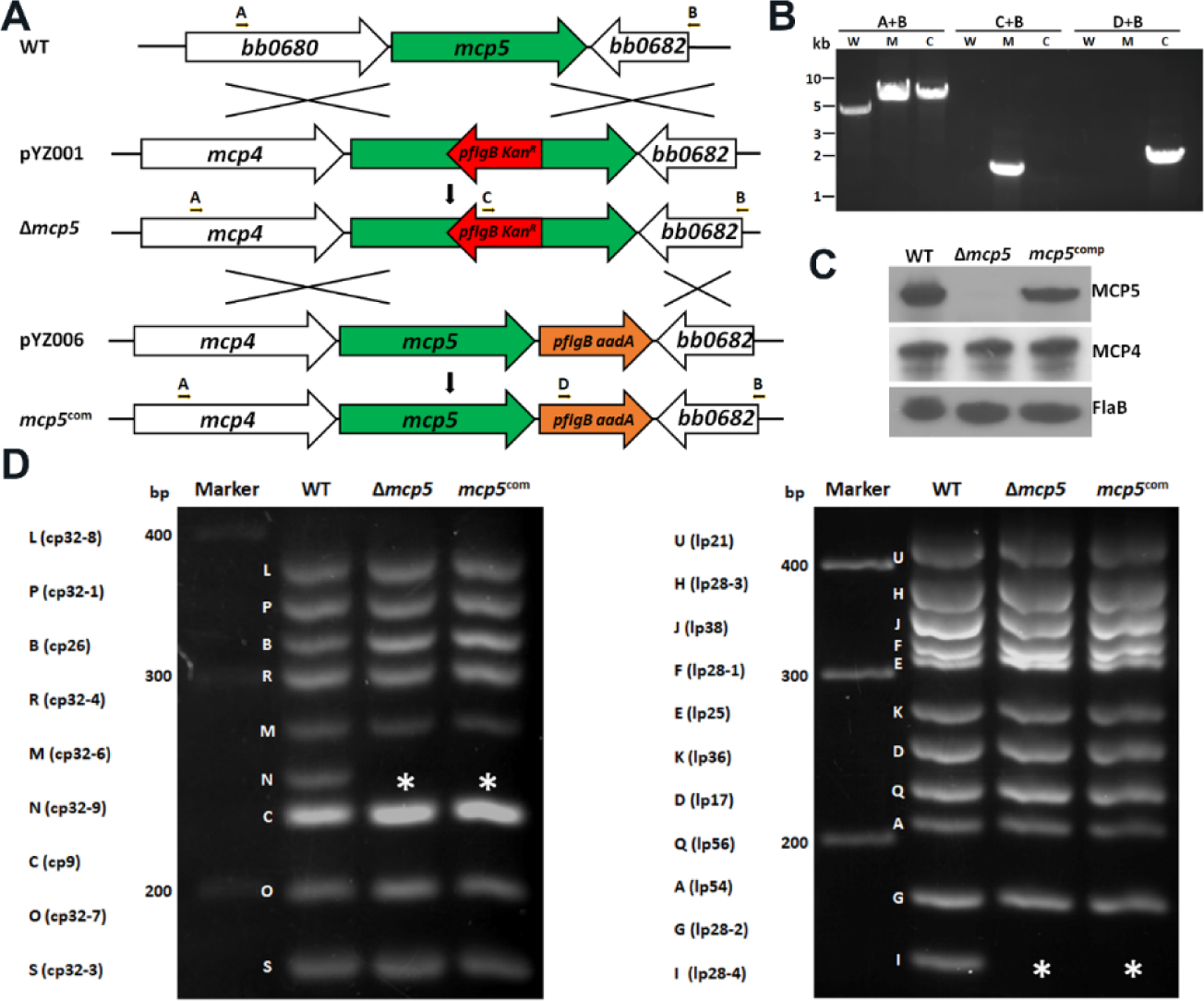
Construction of *mcp5* deletion mutant (Δ*mcp*5) and its isogenic complemented strain (*mcp5*^com^). **(A)** Strategy for constructing the *mcp5* mutant. WT: genomic organization of *mcp5* in *B. burgdorferi* genome. pYZ001: the suicide vector used for deletion of *mcp5*. Arrows indicate the primers used for PCR analyses. pYZ006: the suicide vector used for cis-complementing Δ*mcp*5, generating the complemented strain *mcp5^com^*. The specific primer pairs used are indicated at the top. **(B)** PCR to confirm inactivation and complementation. W, wild-type *B. burgdorferi* strains B31; M, Δ*mcp5*; C, *mcp5*^com^. **(C)** Immunoblotting of MCP4, MCP5 and FlaB levels. *B. burgdorferi* strains B31M (WT), Δ*mcp5* and *mcp5*^com^ strains were cultured in BSK-II medium to stationary phase at 37°C. The whole cell lysates were extracted and were probed with antibodies against MCP5, MCP4, and FlaB (loading control). **(D)** Endogenous plasmid profiles. Endogenous plasmid profiles of each strain by multiplex PCR analyses as previously described [60]. cp, circular plasmid; lp, linear plasmid. Letters on the left indicate the bands corresponding to each endogenous plasmid that was defined previously for the *B. burgdorferi* strain B31 genome [26, 61]. The asterisk (*) indicates the lost plasmids cp-32-6 and lp28-4 respectively.

### MCP5 is required for establishing infection in immune competent C3H/HeN mice

To examine the potential role of MCP5 in the infectious cycle of *B. burgdorferi*, groups of C3H/HeN mice were needle inoculated with WT, Δ*mcp5* and *mcp5*^com^ strains with a dose of 1 × 10^5^ spirochetes/mouse. Ear punch biopsies were collected at 2-, 3-, and 4-weeks post-infection and cultured in BSK-II medium for the presence of spirochetes. At 4-weeks post-infection, all mice were sacrificed and several mouse tissues including ear, joint, heart, skin, and bladder were collected and cultured. Virtually all cultures of tissues from mice inoculated with the wild-type or the complemented strains were positive for *B. burgdorferi* growth. In contrast, only 1 out of 50 cultured mouse tissues showed culture positive from mice infected with the *mcp5* mutant (**Table 1**). To substantiate these observations, spirochetal loads in skin tissues were determined by qPCR. The result showed that the tissues from mice inoculated with the *mcp5* mutant had virtually no detectable or low numbers of spirochetes (**Fig. 7A)**.

**Fig. 7.**
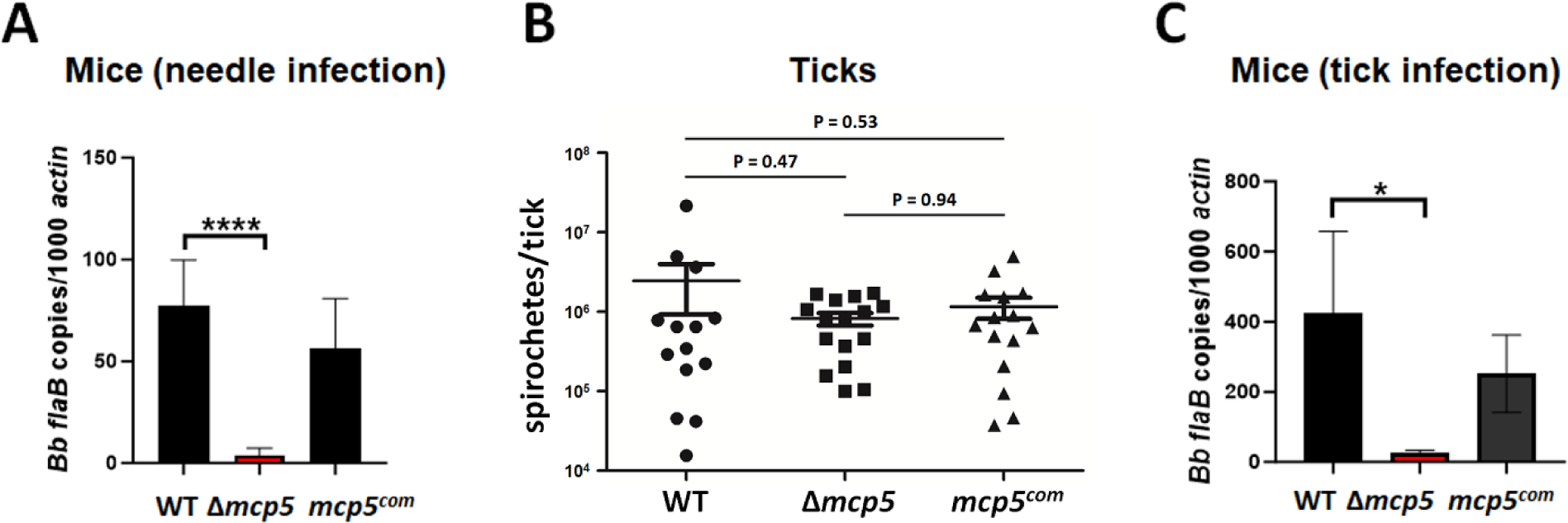
Infection of the *mcp5* mutant in C3H/HeN mice by needle or tick infection. (**A**) Groups of C3H/HeN mice (n = 5) were infected via needle inoculation with 10^5^ cells of WT, Δ*mcp5* and *mcp5^com^*. Mice were euthanized 4 weeks post-infection. DNA were extracted from the skin tissues and number of spirochetes were calculated using qPCR. For qPCR analyses, the copy numbers of *flaB* were normalized to those of the mouse actin gene in each DNA sample. The bar represents the mean values of *flaB* DNA copies calculated from 5 mice. (**B**) Unfed nymphs were artificially infected with WT, Δ*mcp5* and *mcp5^com^* by microinjection, and allowed to feed on naïve mice. Fed nymphs (n=15) collected were lysed and extracted DNA were subjected to qPCR analysis to assess spirochetal loads in ticks. The total copy numbers of *flaB* were reported as per tick DNA sample. The bar represents the mean values of *flaB* DNA copies calculated from 15 fed nymphs. (**C**) Groups of C3H/HeN mice (n = 5) infected by tick bites were euthanized upon tick repletion and subjected to qPCR analyses as described in (**A**). *, *P* < 0.01; ****, *P* < 0.00001 using one-way ANOVA.

**Table 1.** Infection of the *B. burgdorferi mcp5* mutant in immunocompetent mice (C3H/HeN) (4 weeks of post-infection)

| Strain | Dose | Heart | Joint | Ear | Bladder | Spleen | No. of culture positive/No. of tissues* |
| --- | --- | --- | --- | --- | --- | --- | --- |
| B31M | 1x10<sup>5</sup> | 10/10 | 10/10 | 10/10 | 10/10 | 10/10 | 50/50 |
| $\Delta mcp5$ | 1x10<sup>5</sup> | 1/10 | 0/10 | 0/10 | 0/10 | 0/10 | 1/50 |
| *mcp5*<sup>com</sup> | 1x10<sup>5</sup> | 7/10 | 9/10 | 9/10 | 6/10 | 10/10 | 41/50 |

\*Tissues harvested at 4 weeks of post-infection

These data indicate that MCP5 is a virulence factor required for *B. burgdorferi* to establish mammalian infection.

To determine the role of MCP5 in ticks, flat nymphs were artificially infected with wild-type *B. burgdorferi*, the *mcp5* mutant and complemented strains via microinjection [62]. Ticks were then allowed to feed on naïve C3H/HeN mice. Engorged nymphs were collected, and spirochetal loads were assessed by qPCR. The results showed that no significant difference was observed in the estimated spirochetal numbers among ticks harboring each strain (**Fig. 7B)**, suggesting that the *mcp5* mutant is capable of replicating in tick guts during blood meal. To assess the efficiency of transmission from ticks to mice, mouse skin tissues at the site of tick bites immediately upon tick repletion were harvested and subjected to qPCR analyses. The result showed that virtually no or low amounts of *B. burgdorferi* DNA were detected in mouse skin tissues from mice infected with ticks carrying the *mcp5* mutant (**Fig. 7C**), suggesting that although MCP5 is dispensable for replication in ticks, it is required for spirochetes to transmit to the mammalian host.

### The *mcp5* mutant is rapidly cleared in C3H/HeN mice

The inability to establish infection in mice by the *mcp5* mutant could be due to a defect in early colonization or in dissemination. To further investigate the nature of contribution of MCP5 to mammalian infection, immune competent C3H/HeN mice inoculated with wild-type and the *mcp5* mutant were examined at various days post-infection (i.e., day 1, 3, 7 and day 14). At day 1 post-infection, the spirochetal numbers of all strains were similar in the skin tissues of inoculation site (**Fig. 8**). At day 3 post-infection, wild-type spirochetes showed increased numbers at the site of infection and were detected in distal mouse tissues at day 7 and day 14 post-infection. In contrast, at day 3 post-infection, the numbers of *mcp5* mutant at the site of inoculation were significantly reduced (50-fold less than those of the wild-type strain); no mutant spirochetes were detected in distal mouse tissues at day 7 and day 14 post-infection. This data suggests that the *mcp5* mutant spirochetes were quickly cleared at the site of inoculation as early as day 3 post-infection.

**Fig. 8.**
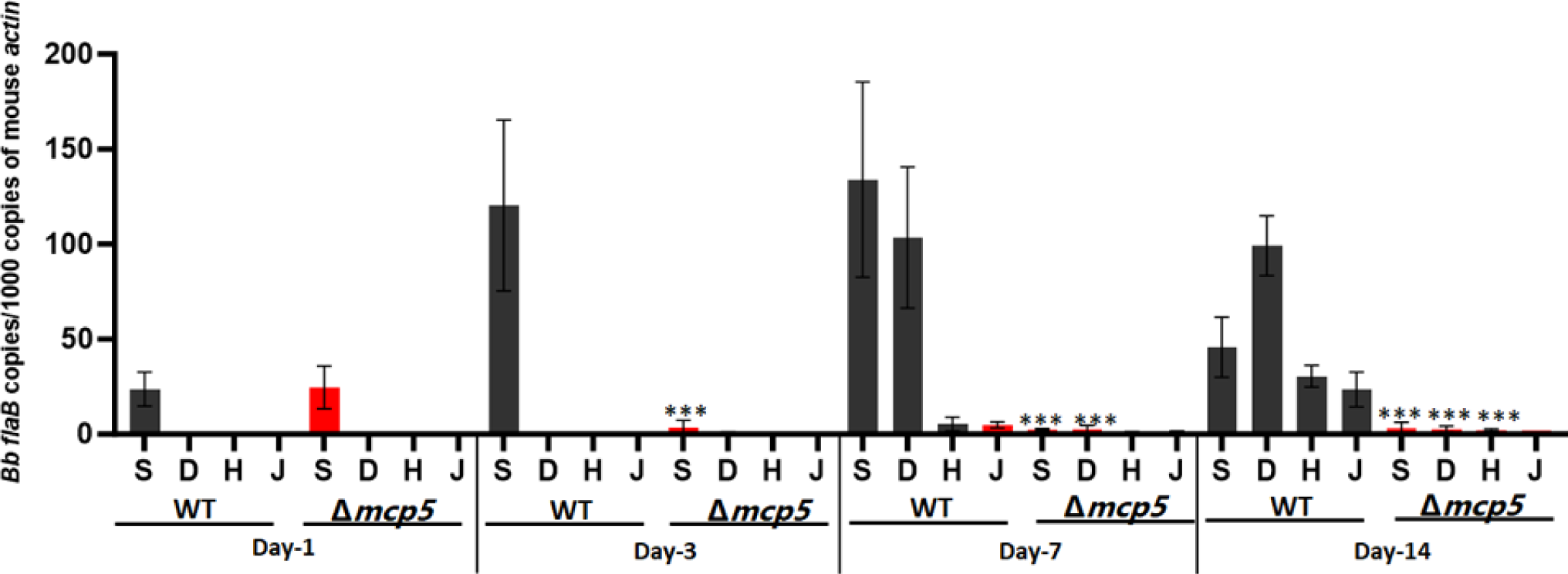
Relative number of *B. burgdorferi* genomes in mouse tissues as determined by qPCR. Groups of C3H/HeN mice (n = 4 per clone per time point) were needle infected with 1×10^5^ WT and Δ*mcp5*, and euthanized after day 1, 3, 7 and 14 days of post-infection. Mouse tissues including skin from the site of infection (**S**), skin at the distant site (**D**), heart (**H**) and joint (**J**) were processed for qPCR analyses. The calculated number of *flaB* copies was normalized to that of mouse actin. The bar represents the mean values of *flaB* DNA copies calculated from each of 4 mice tissues. ***, *P* < 0.0001; using one-way ANOVA.

### The *mcp5* mutant could disseminate and establish infection in <u>N</u>OD <u>S</u>cid <u>G</u>amma (NSG) mice

To gain insight into the essential nature of MCP5 for mammalian infection, immunodeficient SCID mice (deficiency in B and T cells) [63] and NSG mice (NOD-*scid* IL2Rgamma^null^, deficiency in B and T cells, and NK cells, and impaired innate immunity)[64] were needle inoculated with wild-type, the *mcp5* mutant, or the complemented strains. Similar to what was observed in C3H/HeN mice, the *mcp5* mutant failed to infect SCID mice (**Table 2**). Further qPCR analyses showed that the *mcp5* mutant had much lower spirochetal loads than that of the wild-type or complemented strains (**Fig. 9A**). On the other hand, the *mcp5* mutant was able to infect NSG mice that lack adaptive immunity and most of the innate immunity (**Table 2**). qPCR analysis revealed that there were virtually no differences in spirochetal loads between the *mcp5* mutant and wild-type or complemented strains in infected NSG mice (**Fig. 9B**). These results suggest that host innate immunity is responsible for eliminating MCP5-lacking spirochetes, indicating that MCP5 plays an important role in evading host innate immunity.

**Fig. 9.**
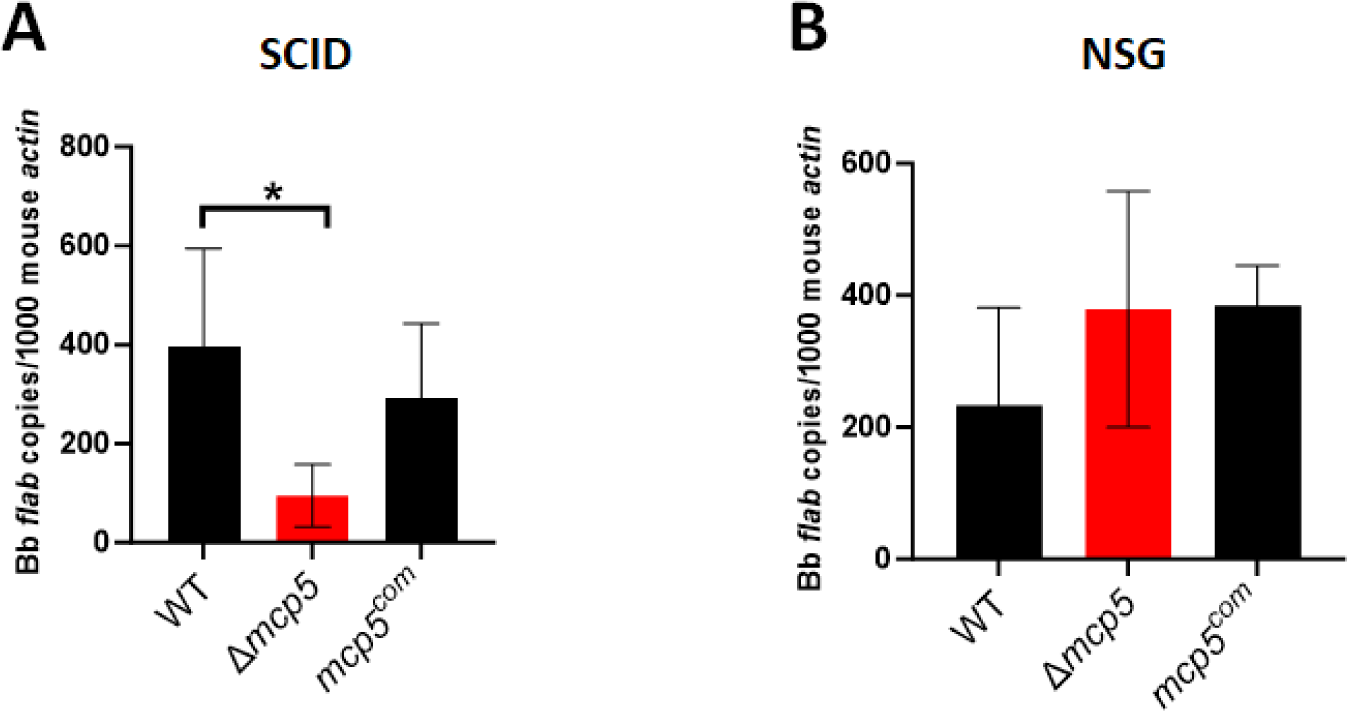
Spirochetal loads in infected SCID and NSG mice. Groups of SCID mice (n = 5 for each data point) (**A**) and NSG mice (n = 5 for each data point) (**B**) were inoculated with 10^5^ of WT, Δ*mcp5*, and *mcp5^com^* strains. Mice were euthanized at 4 weeks post infection and skin (site of infection) were collected and then were subjected to DNA extraction followed by qPCR analyses. The bar represents the mean values of *flaB* DNA copies calculated from 5 infected mice performed in triplicates. *, *p* < 0.01; using one-way ANOVA.

**Table 2.** Infectivity of the *mcp5* mutant in SCID and NSG mice*.

| *Borrelia* Strain | C3H/HeN | SCID Mice | NSG Mice |
| --- | --- | --- | --- |
| **B31M** | 10/10 | 6/6 | 5/5 |
| *Δmcp5* | 1/10 | 1/8 | 7/7 |
| *mcp5<sup>com</sup>* | 7/10 | 5/6 | 6/6 |

\* Determined at 4 weeks of post-infection

## DISCUSSION

The complex nature of the enzootic cycle of *B. burgdorferi* necessitates sensory-guided movement in response to changes in stimuli. In fact, approximately 6% of the *B. burgdorferi* genome contributes to motility and chemotaxis, underscoring their importance in spirochetal complex life cycle [26]. Although much is known about the motility and chemotaxis of *B. burgdorferi*, the knowledge about the chemoreceptors MCPs has been lacking [20, 27]. In this study, we provide evidence that *mcp5* is one of the most abundant and differentially expressed *mcp* genes during the enzootic cycle of *B. burgdorferi*. We further demonstrate that MCP5 is indispensable for *B. burgdorferi* to establish infection in vertebrate hosts, likely by playing an important role in evading host innate immunity.

During the enzootic cycle of *B. burgdorferi*, the Rrp2-RpoN-RpoS pathway is activated during the transmission of spirochetes from ticks to vertebrate hosts and during the infection in vertebrate hosts. This pathway functions as a “gatekeeper” during tick feeding, turning on genes required for spirochetes to establish infection in vertebrate hosts. The inability of the *rrp2*, *rpoN*, *rpoS*, and *bosR* mutants defective in the Rrp2-RpoN-RpoS pathway to express *mcp5* clearly indicates that *mcp5* expression is controlled by Rrp2-RpoN-RpoS during *in vitro* growth conditions (**Fig. 5**). Several lines of evidence suggest that *mcp5* expression is also controlled by Rrp2-RpoN-RpoS during the enzootic cycle of *B. burgdorferi*. Firstly, *mcp5* is differentially expressed during transmission and mammalian infection, correlating with the activation of the Rrp2-RpoN-RpoS pathway. Secondly, MCP5 is required for mammalian infection and for transmission from ticks to mice. How does RpoS control mcp5 expression? Promoter mapping revealed that *mcp5* is transcribed from a σ^70^-type promoter located upstream of *mcp4*, suggesting that *mcp5* is directly regulated by a yet-to-be-identified regulator within the RpoS regulon, or directly by RpoS, given that the promoter sequences between σ^S^ and σ^70^ are nearly indistinguishable. The finding that expression of *mcp5* is also dependent on the Hk1-Rrp1 pathway is quite intriguing, given that unlike Rrp2-RpoN-RpoS, Hk1-Rrp1 is activated in spirochetes replicating in feeding ticks during acquisition and in spirochetes colonizing unfed ticks [33–35]. Interestingly, previous global transcriptomic analyses have also shown that *mcp5* is one of the genes whose expression is affected by both Hk1-Rrp1 and Rrp2-RpoN-RpoS [33, 37, 45, 65–67]. MCP5 appears to be dispensable for replication in the nymphal gut during transmission but is important for spirochetal migration to mice (**Fig. 7**). Whether this is a defect in transmigration from the tick midgut, survival in tick hemolymph, migration to the tick salivary glands, or deposition into mouse dermis, remains to be determined. Given that *mcp5* is regulated by Hk1-Rrp1, it will also be interesting to examine whether MCP5 plays a role in acquisition by ticks and colonization within ticks. Several motility and chemotaxis mutants have been reported to have various phenotypes in ticks. Like the *mcp5* mutant, the *cheY2* mutant can survive in nymphs but fails to transmit *B. burgdorferi* from ticks to mice [68]. A similar phenotype was recently found in the *cheA1* mutant [69]. On the other hand, the *motB* or *cheY3* mutant has reduced spirochetal numbers in feeding ticks [70, 71]. It was speculated that the *motB* or *cheY3* mutant spirochetes could not achieve certain interactions that allows protection against bactericidal factors present in the ingested blood meal in the tick midgut [20].

The infection study in this report revealed that the *mcp5* mutant failed to establish infection and disseminate in both immune-competent and SCID mice but was detectable in all tested tissues in NSG mice (**Table 1 & 2**). SCID mice have a double-strand DNA repair defect that prevents TCR and BCR recombination, resulting in a lack of mature T and B cells, but retaining elements of the innate immune system including natural killer (NK) cells, macrophages, granulocytes and complement proteins [63]. The fact that the *mcp5* mutant cannot infect SCID mice implies that MCP5 may play a role in evading one or more components of the innate immune system that are still functional in SCID mice. NSG mice (NOD.Cg-*Prkdc^scid^ Il2rg^tm1Wjl^*/SzJ) are a more severe immunodeficient strain. They are derived from NOD (Non-Obese Diabetic) mice and carry the *scid* mutation and a targeted mutation in the *IL2rg* gene (interleukin 2 receptor gamma chain). In addition to lacking T cells and B cells, NSG mice also lack functional NK cells and have impaired functions in other innate immune components, including cytokine production and phagocytosis by macrophages, antigen presentation by dendritic cells, and an impaired complement system [64]. The observation that the *mcp5* mutant is capable of infecting NSG mice indicates that the absence of NK cells and impaired other components of innate immunity allow the *mcp5* mutant to establish infection, suggesting that MCP5 is crucial for evading NK cells or other innate immune responses.

How does MCP5 facilitate evasion of the host innate immune response? As a member of the methyl-accepting chemotaxis protein family, MCP5 is predicted to function as a receptor that binds ligands directly or interacts with ligand-binding proteins, transducing signals to downstream signaling proteins to mediate chemotaxis, guiding spirochetes to move toward higher concentrations of attractants or away from repellents [72]. Several chemo-attractants of *B. burgdorferi* have been identified, including serum, glucosamine, N-acetylglucosamine (NAG), glutamate, tick salivary gland, and tick salivary gland protein Salp12 [15, 47–49]. MCP5 is highly expressed *in vitro*; however, to our surprise, the *mcp5* mutant has a normal swimming behavior and still responds to rabbit serum and NAG (**Supplemental Fig. S2**), suggesting that MCP5 is dispensable for chemotaxis *in vitro*. Structural modeling analysis revealed that the N-terminus of MCP5 likely contains a double-Cache (dCache) domain (**Fig. 1**), which comprises a superfamily of the most abundant extracellular sensors in prokaryotes. However, the most N-terminal Cache domain of MCP5, which often binds ligands in related receptors, has relatively low sequence homology to domains of known structure, and accordingly, the 3D prediction does not fully conform to the Cache fold and has quite low confidence (**Fig. 1**). Despite this, we tested several potential ligands of Cache domains, including deoxy- and ribonucleosides (e.g., adenosine, cytidine, uridine, and pyrimidine) and sugars (e.g., D-ribose and pyruvate). There was no significant difference between the wild type and the *mcp5* mutant in sensing these compounds (data not shown). It is highly plausible that MCP5 senses yet-to-be-identified host signals, aiding in the evasion of the innate immune response. Nonetheless, this study demonstrates that MCP5 plays an essential role in the enzootic cycle of *B. burgdorferi*. These findings lay the groundwork for further elucidation of how *B. burgdorferi* utilizes MCP-mediated chemotaxis and motility to navigate between and within the tick vector and the vertebrate host.

## MATERIALS AND METHODS

### Ethics statement

All animal experiments were approved by the IACUC committee of Indiana University School of Medicine under protocol number # 20126. All experiments were in accordance with the institutional guidelines.

### *B. burgdorferi* strains and culture conditions

Low-passage, virulent *B. burgdorferi* strain B31 was used in this study [73]. Spirochetes were cultivated in Barbour-Stoenner-Kelly (BSK-II) medium supplemented with 6% rabbit serum (Pel-Freez Biologicals, Rogers, AR) [79] at 37°C with 5% CO_2_. At the time of growth, appropriate antibiotics were added to the cultures with the following final concentrations: 300 μg/ml for kanamycin and 50 μg/ml for streptomycin. The constructed suicide vectors for inactivation (pYZ001) and complementation (pYZ006) were maintained in *Escherichia coli* strain DH5α. The antibiotic concentrations used for *E. coli* selection were as follows: kanamycin (50 μg/ml) and streptomycin (50 μg/ml). A list of all the *B. burgdorferi* strains and plasmids used in the present study are represented in supplemental material (see Table S1).

### Immunoblot analysis

Spirochetes from various stages of growth were harvested by centrifuging at 8,000 × g for 10 min, followed by three times washing with PBS (pH 7.4) at 4°C. Cell pellets were suspended in SDS buffer containing 50 mM Tris-HCl (pH 8.0), 0.3% sodium dodecyl sulfate (SDS) and 10 mM dithiothreitol (DTT). Cell lysates (10^8^ cells per lane) were separated by 12% SDS-polyacrylamide gel electrophoresis (PAGE) and transferred to nitrocellulose membranes (GE-Healthcare, Milwaukee, WI). Membranes were blotted with rat polyclonal antibody against MCP5 (1:3,000 dilution) and monoclonal antibody against FlaB (1:1,000 dilution), followed by goat anti-rat lgG-HRP secondary antibody (1:1,000; Santa Cruz Biotechnology). Detection of horseradish peroxidase activity was determined using the enhanced chemiluminescence method (Thermo Pierce ECL Western Blotting Substrate) with subsequent exposure to X-ray films.

### AlphaFold model generation and analysis

To build models for *B. burgdorferi* (Bb) MCPs, protein sequences for *B. burgdorferi* MCP1 (O51525), MCP2 (O51542), MCP3 (O51543), MCP4 (O51623) and MCP5 (O51624) were retrieved from the UniProt database and submitted for automated model-building using AlphaFold2 [51]. All parameters were kept at their default values for model building.

### Generation of *mcp5* deletion mutant and its isogenic complemented strain

To inactivate *mcp5* in *B. burgdorferi* strain B31, a suicide vector pYZ001 was constructed. The regions of DNA corresponding to 1.5 kb upstream and downstream of *mcp5* were PCR amplified using specific set of primers PRYZ001/PRYZ002 (up) and PRYZ003/PRYZ004 (downstream) from *B. burgdorferi* (see Table S2). The PCR products were cloned into a suicide plasmid pUC-Kan containing a *flaB* promoter driven kanamycin marker (*kan*) at ApaI and XbaI restriction sites (for upstream fragment) and HindIII and BamHI sites (for downstream fragment), respectively. The resulting suicidal plasmid pYZ001 was transformed into wild-type *B. burgdorferi* B31 as previously reported [62], and positive clones were selected based on kanamycin resistance and further validated by PCR and Western blot analyses. Endogenous plasmid profiles were performed for the *mcp5* mutant clones as previously described [60, 74].

For *cis* complementation (gene replacement), the suicidal plasmid pYZ006 was generated as follows. The regions containing the full length *mcp5* were PCR amplified using specific sets of primers PRYZ009/PRYZ010 (upstream fragment) and PRYZ011/PRYZ012 (downstream fragment) from *B. burgdorferi* genomic DNA (see Table S2 in the supplemental material). The upstream fragments were then cloned into the suicide vector pCT007 at ApaI and SalI restriction sites and XmaI and XbaI sites, respectively. The resulting suicidal plasmid pYZ006 was transformed into the *mcp5* mutant, and streptomycin resistant and kanamycin sensitive clones were selected, confirmed by PCR and Western blot analyses.

### 5′ Rapid amplification of cDNA end (5′ RACE) analysis

This assay was conducted as previously described [75]. In brief, wild-type *B. burgdorferi* B31 A3-68 cells were cultivated at 37°/pH 7.5 until late log phase and then harvested for RNA extraction using NucleoSpin RNA kit, following the manufacturer’s instruction (Macherey-Nagel, Bethlehem, PA). 5′ RACE was carried out using SMARTer RACE 5′/3′ Kit (Takara Bio USA, Mountain View, CA) to identify the transcription start site (TSS) of *mcp4* and *mcp5* genes following the manufacturer’s protocol. Primers used for the 5’RACE (Primers) were listed in Table S2.

### *B. burgdorferi* motility and chemotaxis assays

Bacterial cell motility (wild-type *B. burgdorferi* B31M, the *mcp5* mutant, the *mcp5* complemented strain) was measured using a computer-based motion tracking system as previously described [47]. The *flaB* mutant (*flaB^-^*), a previously constructed non-motile mutant [76], served as a negative control. Briefly, late-log phase *B. burgdorferi* cultures were first diluted (1:1) in BSK-II medium and then 10 μl of the diluted cultures were mixed with an equal volume of 2% methylcellulose, and then subjected to dark-field microscopy. Spirochetes were video captured with iMovie software on a Mac computer and then exported as QuickTime movies, which were further imported into OpenLab (Improvision Inc., Coventry, UK) where the frames were cropped, calibrated, and saved as LIFF files. The software package Velocity (Improvision Inc.) was used to track individual moving cells to measure their velocities. For each bacterial strain, at least 20 cells were recorded for up to 30 sec. The average cell swimming velocities (μm/s) of tracked cells were calculated. Swimming plate assays were performed using 0.35% agarose with BSK-II medium diluted 1:10 with Dulbecco’s phosphate-buffered saline (DPBS, pH 7.5), as previously described [22, 23, 76]. The plates were incubated for 4–5 days at 34°C in the presence of 3.4% CO_2_. The diameters of swim rings were measured and recorded in millimeters (mm). The average diameters of each strain were calculated from four independent plates. Capillary tube assays were carried out as previously documented with minor modifications [47]. In brief, *B. burgdorferi* cells were grown to late-log phase (∼5–7 × 10^7^ cells/ml) and harvested by low-speed centrifugations (1,800 × *g*). The harvested cells were then resuspended in the motility buffer. Capillary tubes filled with either the attractant (0.1 M N-acetylglucosamine dissolved in the motility buffer, or 0.5% rabbit serum) or only motility buffer (negative control) were sealed and inserted into microcentrifuge tubes containing 200 μl of resuspended cells (7 × 10^8^ cells/ml). After 2 hrs incubation at 34°C, the solutions were expelled from the capillary tubes, and spirochete cells were enumerated using Petroff-Hausser counting chambers under a dark-field microscope. A positive chemotactic response was defined as at least twice as many cells entering the attractant-filled tubes as the buffer-filled tubes. For the tracking, swimming plate, and capillary assays, the results are expressed as means ± standard errors of the means (SEM). The significance of the difference between different strains was evaluated with an unpaired Student *t* test (*P* value < 0.01).

### Mouse infection studies

Four-week-old C3H/HeN mice, C3H/SCID and NSG mice (Harlan, Indianapolis, IN) were subcutaneously inoculated with two doses of spirochetes (1×10^5^ and 1×10^6^) respectively. Ear punch biopsy samples were taken at 2- and 3-weeks post-injection. At 4 weeks post-injection, mice were euthanized, and multiple tissues (i.e., ear, joint, heart, skin and bladder tissues from each mouse) were harvested. All tissues were cultivated in 2 ml of the BSK-II medium (Sigma-Aldrich, St. Louis, MO) containing an antibiotic mixture of phosphomycin (2 mg/ml), rifampin (5 mg/ml), and amphotericin B (250 mg/ml) (Sigma-Aldrich) to inhibit bacterial and fungal contamination. All cultures were maintained at 37°C and examined for the presence of spirochetes by dark-field microscopy beginning from 5 days after inoculation. A single growth-positive culture was used as the criterion to determine positive mouse infection.

### qRT-PCR analyses

For identifying the expression pattern of *mcp5 in vitro,* wild-type *B*. *burgdorferi* strain B31 was cultured in BSK-II medium at various conditions. RNA samples were extracted from *B. burgdorferi* cultures using the RNeasy mini kit (Qiagen, Vanelcia, CA) according to the manufacturer’s protocols, followed by an on-column Digestion RNase-free DNase I treatment (Promega, Madison, WI). Quality of the isolated RNA was confirmed using PCR amplification of *B. burgdorferi flaB* (to check for DNA contamination). The cDNA was synthesized using the SuperScript III reverse transcriptase with random primers (Invitrogen, Carlsbad, CA). Quantitative PCR (qPCR) was performed in triplicate on an QuantStudio™ 3 thermocycler. Calculations of relative levels of transcript were normalized with *flaB* transcript levels as previous reported[77].

For quantifying *mcp5* and *ospC* expression in infected mice, four-week-old C3H/HeN mice were injected with wild-type *B*. *burgdorferi* strain B31M at a dose of 1×10^4^ spirochetes per mouse. Mice were euthanized at different time points as indicated and mouse tissues were harvested and homogenized using the FastPrep-24 (MP Biomedicals). Total RNA was isolated using the TRIzol reagent (Thermo Fisher Scientific) according to the manufacturer’s instructions. To eliminate DNA contamination, samples were further digested with RNase-free DNaseI (Qiagen), purified using the RNeasy mini kit (Qiagen) and analyzed with NanoDrop Spectrophotometer (Thermo Fisher Scientific). cDNA was synthesized using the PrimeScript 1st strand cDNA Synthesis Kit (Takara Bio USA). For RNA analysis of spirochetes in ticks, 10 groups of fed larvae (3 ticks per group), 3 groups of unfed nymphs (40 ticks per group), and 10 groups of fed nymphs (one tick per data point) were used. Given the low levels of bacterial RNA in mouse tissues, the specific primers for each gene target were used for cDNA synthesis instead of random primers. To quantify the transcript levels of genes of interest, an absolute quantitation method was used to create a standard curve for the qRT-PCR assay according to the manufacturer’s protocol (Strategene, La Jolla, CA). Briefly, the PCR product of the *flaB* gene served as a standard template. A series of tenfold dilutions (10^2^-10^7^copies/ml) of the standard template was prepared, and qRT-PCR was performed to generate a standard curve by plotting the initial template quantity against the Ct values for the standards. The quantity of the targeted genes in the cDNA samples was calculated using their Ct values and the standard curve. The samples were assayed in triplicate using the ABI 7000 Sequence Detection System and PowerUp SYBR Green Master Mix (Applied Biosystems). The levels of the target gene transcript were reported as per 1000 copies of *flaB*. The primers used for the qRT-PCR analysis are depicted in Table S2.

### Microinjection of *B. burgdorferi* into nymphal ticks

Microinjection and the tick-mouse experiments were approved by the IACUC committee of Indiana University School of Medicine under protocol number #11339. *I. scapularis* nymphs were obtained from the Tick Rearing Facility at Oklahoma State University (Stillwater, OK). Microinjection was used to introduce spirochetes into the gut of nymphs as previously described [62]. Briefly, each *B. burgdorferi* strain was cultivated under normal conditions in BSK-II medium in the presence of corresponding selective antibiotics. Spirochetes were harvested by centrifugation and concentrated in BSK-II to a density of 5 × 10^8^ spirochetes/ml. A total of 10 μl of the cell suspension was then loaded into a 1mm diameter glass capillary needle (World Precision Instruments Inc.) by using a micro loader (Eppendorf AG). The bacterial suspension was then injected into the rectal aperture of unfed nymphal ticks by using a FemtoJet microinjector system (Eppendorf AG) as previously described [62].

### Assessment of spirochete transmission to mice by encapsulated nymphs

Transmission of spirochetes from *I. scapularis* ticks to C3H/HeN mice was assessed using artificially infected nymphs via microinjection as described above. Mice were anesthetized, infected ticks were confined to a capsule affixed to the shaved back of a naive C3H/HeN mouse (9 ticks per mouse). The ticks were allowed to feed to repletion (3 to 5 days) and then collected for DNA extraction. Subsequently, each sample of tick DNA was used to determine bacterial burdens by qPCR. Infected mice were then subjected to qPCR analysis to assess spirochetal burden in mouse tissues or culturing for *Borrelia* growth.

### Extraction of tick DNA

DNA was isolated from engorged nymphs using the DNeasy® Blood & Tissue Kits (QIAGEN) according to the manufacturer’s instructions. Spirochete burdens within infected ticks were assessed with primer pairs of q-*flaB*-F/R and q-*Tactin*-F/R (see Table S2 in the supplemental material). Absolute copy numbers of *flaB* are quantified as spirochete loads in ticks.

## ACKNOWLEDGEMENT

We express our gratitude to Dr. Zhiming Ouyang for generously supplying the *bosR* mutant strain. Funding for this research was partly supported by NIH grants AI083640 and AI152235 (to X. F. Yang), R35GM122535 and AI148844 (to B. Crane), AI078958 (to C. Li), and National Natural Science Foundation of China 82072310 (to Y. Lou). Additionally, we acknowledge the use of facilities supported by the research facilities improvement program grant number C06 RR015481-01 from the National Center for Research Resources, NIH.

